# Co-sedimentation is the key to the structural investigation of wild-type FAT10

**DOI:** 10.64898/2026.02.06.704312

**Authors:** Charlotte Weiss, Barbara Perrone, Nicola Catone, Annette Aichem, Guinevere Mathies

**Affiliations:** Department of Chemistry, University of Konstanz, Konstanz 78464, Germany; Bruker Switzerland, Fällanden 8117, Switzerland; Institute of Cell Biology and Immunology Thurgau, Kreuzlingen 8280, Switzerland

## Abstract

Under inflammatory conditions, the ubiquitin-like modifier FAT10 serves as a tag for protein degradation by the 26S proteasome. FAT10 is degraded along with its substrates and this process is independent of the segregase VCP/p97, which, in the regular ubiquitin pathway of degradation, is required if a substrate lacks a disordered initiation region. FAT10 itself is loosely folded and its tendency to aggregate has complicated investigations of its structure, interaction, and function. Recently hydrogen-deuterium exchange in combination with mass spectrometry has suggested that, in preparation of degradation by the proteasome, the adapter protein NUB1 traps FAT10 in a mostly unfolded state by capturing a β-strand. β-strand capture was subsequently confirmed by magic-angle spinning (MAS) NMR spectroscopy of a stabilized variant of the N-domain of FAT10 in complex with NUB1L, the longer splice variant of NUB1. MAS NMR, in addition, revealed that the N-domain of FAT10 and NUB1L form a fuzzy complex and that the N-terminus of FAT10 is positioned for initiation of degradation by specific non-covalent interaction with NUB1L. Here, we report the investigation of the wild-type N-domain of FAT10 by MAS NMR. Co-sedimentation with NUB1L yields high-quality spectra, which enable sequential assignment of resonances. Through the lens of MAS NMR, the complexes of the wild-type and stabilized N-domain of FAT10 with NUB1L are identical. The N-terminus of FAT10 again shows up prominently in the spectra, even though the residue is this time an Ala, not a Gly. Our experience suggests that co-sedimentation in combination with MAS NMR is generally helpful in the exploration of conditional folds of intrinsically disordered proteins.

## INTRODUCTION

Intrinsically disordered proteins (IDPs) and intrinsically disordered protein regions (IDRs) do not have a single, well-defined equilibrium structure. Instead they exist as dynamic, structurally heterogeneous ensembles of conformers.^1,2^ Their biological functions directly relate to these ensembles, which are strongly influenced by local conditions.^3–5^ Interaction with binding partners occurs via molecular recognition and binding-induced folding, i.e., the IDP undergoes a transition from disorder to order.^2^ The ordering upon binding is, however, often not complete, giving rise to so-called fuzzy complexes.^6^ AlphaFold^7^ is remarkably good at predicting conditional folds of IDPs,^8^ but cannot provide information about ensembles of conformers and, consequently, struggles with fuzzy complexes. Experimental investigation of IDPs and their interactions thus remains pertinent.

Nuclear magnetic resonance (NMR) spectroscopy has provided a wealth of information about the structure and dynamics of IDPs in solution.^9^ Tailored pulse sequences enable sequential assignment of residues, in spite of the reduced spread of ^1^H resonances.^10^ Nuclear spin relaxation rates provide information about local dynamics^11^ and paramagnetic relaxation enhancements report on long-range interactions and conformational ensembles.^12^ These experiments have also been of great value in the investigation of the interactions of IDPs with other proteins, for example in chaperone-client complexes.^13^

For the characterization of IDPs in the condensed state, magic-angle spinning (MAS) NMR is well suited. Sequential assignments are made based on multi-dimensional ^13^C or, with the advent of ultrafast MAS, ^1^H detected spectra.^14,15^ The choice of experiments depends on particular conditions such as access to instrumentation, available amount and size of the protein, and structural homogeneity in the sample. Subsequent determination of protein structure requires angle and distance constraints. The former follow from analysis of chemical shifts, while the latter require measurements of dipolar couplings in dedicated experiments on, in the case of ^13^C detection, a specifically-labeled sample. In this way, the structures of amyloid fibrils have been solved at atomic resolution. Examples include Aβ^16–18^, α-synuclein^19,20^, and β_2_-microglobulin^21^. In line with expectations for IDPs, amyloid fibrils tend to have a fuzzy coating^22^ (i.e., the ordering is not complete), which can in some cases be observed with solution-state NMR experiments.

The small protein FAT10 (human leukocyte antigen F adjacent transcript 10) targets substrates for degradation by the 26S proteasome.^23–25^ The FAT10 pathway is active under inflammatory conditions and bears resemblance to the canonical ubiquitin pathway for proteasomal degradation.^26^ Both FAT10 and ubiquitin become covalently attached to substrates via an enzyme cascade consisting of an activating enzyme (E1), a conjugating enzyme (E2), and a ligase (E3).^27,28^ A polyubiquitin chain is required for substrate degradation^29^ and FAT10 consists of two ubiquitin-like domains, the N- and C-domain, which are connected by a flexible linker.^30,31^ There are also notable differences. If a ubiquitin-labeled substrate is tightly folded, the segregase VCP (valosin containing protein or p97) initiates degradation by unfolding a ubiquitin,^32^ which is otherwise left intact and reused in the cell. Degradation of FAT10-labeled substrates is VCP-independent^31^, but is only significant in the presence of adapter protein NUB1L (NEDD8-ultimate buster 1 long),^33,34^ which is known to interact with the N-domain of FAT10^35^ and with the ubiquitin receptors RPN1 and RPN10 of the proteasome^36^. Also, FAT10 is degraded along with its substrates^23^.

FAT10 is poorly soluble and has a tendency to aggregate.^37,38^ Its melting temperature is with 41 °C relatively low, compared to 83 °C for ubiquitin.^31^ Molecular dynamics simulations have highlighted the absence of long-range salt bridges in FAT10 and revealed low resistance to mechanical unfolding.^39^ These properties have interfered with investigations of structure, interaction, and function. Only by deleting the first seven residues, Theng *et al*. were able to sufficiently stabilize the N-domain for structure determination by solution-state NMR.^40^ Residues P59-T73 were not visible in HSQC (heteronuclear single quantum coherence) spectra and this observation, together with an absence of slowly exchanging amide protons, let the authors to conclude that the N-domain experiences global folding and unfolding. Aichem *et al*. determined the structure of a stabilized, Cys-free mutant (C7T, C9T, C134L, C162S) of FAT10 by X-ray diffraction (for the N-domain) and solution-state NMR (for the C-domain).^31^ Both domains show the characteristic β-grasp fold of ubiquitin-like modifiers^41^, but their surfaces differ from ubiquitin and each other. Intriguingly, the stabilization of FAT10 by mutation of the Cys residues had no effect on substrate conjugation, but degradation rates were clearly reduced.^31^

Recently, Arkinson *et al*. used hydrogen-deuterium exchange detected by mass spectrometry to investigate the conformation and solvent accessibility of wild-type FAT10.^42^ Peptides from both the N- and the C-domain show a bimodal distribution, confirming the coexistence of folded and unfolded states. In the presence of NUB1 (the shorter splice variant of NUB1L), peptides throughout the N-domain are exposed, with exception of the β5-strand. This observation was in agreement with a structure predicted by AlphaFold-Multimer^43^ in which the N-domain of FAT10 is partially unfolded and residues of the former β5-strand are engaged in an intermolecular, anti-parallel β-sheet with NUB1L. Arkinson *et al*. concluded that, in preparation of proteasomal degradation, NUB1 traps the N-domain of FAT10 in an unfolded state by β-strand capture. Using a combination of methods including cryo-electron microscopy, they could additionally show that binding of FAT10 induces an open conformation of NUB1, which allows its ubiquitin-like domain to interact with the RPN1 subunit of the regulatory particle. Unfortunately, resolution was not sufficient to pinpoint the interaction between FAT10 and NUB1.

Meanwhile our group performed MAS NMR on the stabilized, Cys-free N-domain of FAT10 (N-FAT10-C0), both isolated and in complex with NUB1L.^44^ Sequential assignment of amino acid residues followed by empirical prediction of torsion angles and secondary structure confirmed that isolated N-FAT10-C0 is in the ubiquitin-like β-grasp fold. In addition to the flexible start and end tails, a stretch of residues leading up to and partially covering the β5-strand (I68-L76) were missing from the spectra. We attributed this to disorder arising from local structural frustration, suggesting that this stretch of residues is primed for β-strand capture. To prepare the complex of N-FAT10-C0 and NUB1L we used co-sedimentation by ultracentrifugation^45,46^ of an equimolar solution directly into the MAS rotor. Spectra of the complex show the selective survival of a small set of resonances, which we could sequentially assign to I68-V81, W17, and the N-terminus (G4) and to an additional eleven residues for which the spectra contained no information about the position in the sequence. Torsion angle and secondary structure prediction classified T73-V81 as β-strand, thus providing atomic-level experimental evidence for the existence of the intermolecular β-sheet AlphaFold-Multimer had predicted. Further secondary structure elements of N-FAT10-C0 predicted by AlphaFold-Multimer with high confidence were notably absent from the MAS NMR spectra. We concluded that these parts of N-FAT10-C0 are, in reality, disordered. The discrepancy likely arises from the tendency of AlphaFold to stabilize conditionally folded elements^8^ and, following the MAS NMR experimental data, we proposed the formation of a fuzzy complex: NUB1L acts as a holdase for N-FAT10-C0 and positions the N-terminus to initiate the degradation by the proteasome. Participation of the observed resonances in the intermolecular interface between N-FAT10-C0 and NUB1L was confirmed with a series of double-REDOR (Rotational-Echo Double-Resonance) filtered^47^ experiments.

Here, we report the investigation of the N-domain of wild-type FAT10 (N-FAT10-WT) and its interaction with NUB1L by MAS NMR. Increased β-sheet content confirms aggregation of isolated N-FAT10-WT upon concentration and sedimentation in the MAS rotor. Co-sedimentation by ultracentrifugation is again a successful strategy for preparation of the complex of N-FAT10-WT and NUB1L if care is taken to keep the concentration of N-FAT10-WT at 1 mg/L. The resolution of resulting ^13^C-^13^C and ^15^N-^13^C correlation spectra is very good. For the purpose of sequential assignment, we record ^15^N-^13^C-^13^C correlation spectra with a Bruker MAS CryoProbe^48^, which offers, as a result of cooling of its coil and electronics, a 3-4 fold higher sensitivity compared to regular MAS NMR probes. Somewhat to our surprise as well as reassurance, spectra show that the complexes of N-FAT10-WT and N-FAT10-C0 with NUB1L are identical. Even though the residue is now an Ala (A5), not a Gly, we observe a pronounced signal form the N-terminus, indicating that also in the complex with N-FAT10-WT it has a regular, stable structure. Our experience suggests that co-sedimentation in combination with MAS NMR is generally helpful in the exploration of the conditional folding of IDPs, for example upon interaction with a binding partner.

## MATERIALS and METHODS

### Expression and Purification of U-^13^C,^15^N-N-FAT10-WT

Wild-type N-domain of human FAT10 (amino acids 2-86) and an N-terminally truncated variant thereof (amino acids 5-86) were cloned into pSUMO vectors for expression as His_6_-SUMO-fusion proteins. Expression was performed in *E. coli* BL21-CodonPlus(DE3)-RIPL competent cells (Agilent Technologies). For uniform ^13^C and ^15^N labelling (U-^13^C,^15^N), cells were grown in M9 minimal medium supplemented with U-^13^C_6_-D-glucose and ^15^N-ammonium chloride. The purification protocol is analogous to the protocol for the Cys-free N-domain (N-FAT10-C0)^31,44^ and is detailed in the Supporting Information, where SDS-PAGE analysis is also provided (Figure S1). Briefly, bacterial cells were lysed and the supernatant was applied to nickel affinity chromatography. After buffer exchange, Ulp1 protease was added for cleavage of the purification tag. The Ulp1 protease recognizes the three-dimensional fold of the SUMO-tag preserving the sequence of N-FAT10, which starts, for both variants, with an alanine residue. The His-tagged protease and respective cleavage byproducts were separated by a second nickel affinity chromatography. Further purification was achieved by size-exclusion chromatography yielding up to 3 mg per litre of M9 medium.

### Expression and Purification of NUB1L

As described in our previous work,^44^ human NUB1L (amino acids 2-615) was also expressed as a His_6_-SUMO-fusion protein in *E. coli* BL21CodonPlus(DE3)-RIPL competent cells. The pSUMO plasmid was kindly provided by the Institute of Cell Biology and Immunology Thurgau. To produce natural abundance NUB1L, cells were grown in LB medium. Purification required the same steps as for U-^13^C,^15^N-N-FAT10-WT. Yield was up to 15 mg per litre of LB medium.

### Sample packing for MAS NMR

Ultracentrifugation (Beckman Coulter Optima L90K, SW 41 Ti Swinging-Bucket Rotor) was used to ensure dense protein packing. For measurements at 18.8 T, with a standard E-free HCN Bruker MAS probe, we used a home-made tool that enabled direct packing into a regular-wall zirconia 3.2 mm MAS rotor (maximum sample volume 32 µL). The tool can hold 1 mL of protein solution. In contrast to N-FAT10-C0, for which concentrations up to 25 mg/mL can be reached and crystallization is possible, isolated N-FAT10-WT loses its solubility and starts to precipitate at concentrations above 1-2 mg/mL. Nevertheless, for the purpose of characterization by MAS NMR, a suspension of U-^13^C,^15^N-N-FAT10-WT (amino acids 2-86) was ultracentrifuged for 4 h at RCF 210000 g and 4 °C. After removal of the supernatant, the MAS rotor, containing a white precipitate, was closed with a Vespel cap.

To form a complex of N-FAT10-WT and NUB1L, co-sedimentation by ultracentrifugation was used. 3 mL of U-^13^C,^15^N-N-FAT10-WT (amino acids 5-86) at a concentration of 1 mg/mL was mixed with 1 mL of natural abundance NUB1L with a concentration of 20 mg/mL, leading to a 1:1 molar ratio. The packing tool containing this solution was centrifuged at RCF 210000 g and 4 °C. Over the course of several days, the run was interrupted repeatedly to remove supernatant and gradually add the original mixture in full. Packing efficiency was assessed by UV-Vis absorbance measurements of the supernatant at 280 nm. The MAS rotor was closed with a Vespel cap.

A tool to safely pack, by ultracentrifugation, a sample into a thin-wall silicon nitride 3.2 mm CryoProbe MAS rotor (maximum sample volume 87 μL)^48^ is not available. Hence, for measurements at 14.1 T with a Bruker MAS CryoProbe, a solution of U-^13^C,^15^N-N-FAT10-WT (amino acids 5-86) and natural abundance NUB1L as described above was first centrifuged in a polypropylene ultracentrifuge tube for several days at RCF 210000 g and 4 °C. After removal of the supernatant, the co-sediment (pellet) was transferred into a CryoProbe MAS rotor with a swinging-bucket benchtop centrifuge (2 h at RCF 3100 g and 4 °C). Teflon spacers were placed at the top and bottom of the CryoProbe MAS rotor, giving a reduced sample volume of 47 µL.

### MAS NMR Spectroscopy

A first series of experiments was performed at 18.8 T (^1^H Larmor frequency of 800.3 MHz) on a Bruker Avance NEO spectrometer equipped with a 3.2 mm E-free HCN Bruker MAS probe. The spinning frequency was 14.5 kHz and the sample temperature was 4 °C. Sample integrity was verified by ^1^H-^13^C and ^1^H-^15^N cross-polarization (CP) spectra as well as ^13^C-^13^C correlation spectra recorded with the dipolar assisted rotational resonance (DARR)^49^ pulse sequence. A mixing time of 10.0 ms is suitable for the observation of predominantly one- and two-bond contacts. For N-FAT10-WT in complex with NUB1L, a ^15^N-^13^C correlation spectrum was recorded as well with the z-filtered transferred echo double-resonance (ZF TEDOR)^50^ pulse sequence. A mixing time of ∼1 ms is suitable to detect N-C_α_ and N-C’ magnetization transfers along the protein backbone and, in the sidechains, to carbon nuclei directly bonded to nitrogen nuclei.

A second series of experiments was performed at 14.1 T (^1^H Larmor frequency of 600.3 MHz), again on a Bruker Avance NEO spectrometer, this time equipped with a Bruker 3.2 mm MAS CryoProbe^48^. The spinning frequency was 12.0 kHz and the sample temperature was 4 °C. After verification of sample integrity, ^15^N-^13^C correlation spectra were recorded using SPECIFIC-CP^51^ to transfer magnetization from backbone nitrogen nuclei either to backbone C’ or C_α_ nuclei. For the purpose of sequential assignment of N-FAT10-WT in complex with NUB1L, ^15^N-^13^C-^13^C NCOCX and NCACX spectra were recorded with SPECIFIC-CP and COmbined 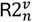 (CORD)^52^ homonuclear carbon mixing. Non-uniform sampling of 42% was used, per its implementation in TopSpin.^53^

An overview of the MAS NMR experiments and the acquisition parameters are given in Tables S1 and S2. Spectra were processed in NMRPipe^54^. NCOCX and NCACX spectra were reconstructed with the Iterative Soft Thresholding (IST) method^55^ as implemented in NMRPipe. The CcpNmr Analysis V3 software was used for the assignment of protein resonances.^56^

### Torsion Angle and Secondary Structure Prediction

Backbone torsion angles Φ and Ψ and secondary structure were predicted empirically based on chemical shifts of assigned N, C’, C_α_ and C_β_ nuclei.^57,58^. For this purpose, we used the program TALOS-N,^59^ which combines a set of trained neural networks with efficient mining of a database of proteins of which both the X-ray structure and the chemical shifts are known.

### AlphaFold Modelling

AlphaFold^7^ v2.3.2 was downloaded from Github (https://github.com/google-deepmind/alphafold/tree/v2.3.2) and installed on the local Scientific Compute Cluster of the University of Konstanz (SCCKN). To run AlphaFold in multimer mode,^43^ a multi-sequence FASTA file including N-FAT10-WT (amino acids 5-86) and NUB1L (amino acids 1-615) was provided. Database presets were set to reduced and the maximum template release date was 2004-11-28. Five seeds were run per model, resulting in 25 structure predictions. These were ranked by model confidence using a weighted combination of predicted template modelling (pTM) and Interface pTM (ipTM) scores (model confidence = 0.8·ipTM + 0.2·pTM). Only the best ranked model was relaxed. Structure visualization was performed with UCSF ChimeraX.^60^

## RESULTS

The ^1^H-^13^C CP spectrum of isolated N-FAT10-WT, packed densely into a MAS rotor, is shown in Figure 1a (chartreuse). Resolution is poor, particularly in the C_α_ region (50-68 ppm) and in the carbonyl region (170-185 ppm), which shows scarcely any features, except for a foot from Glu C_δ_. At the edges of the aliphatic region, characteristic narrow lines from Thr C_α_/C_β_ and Ile C_δ1_ nuclei in well-defined conformations are absent. The backbone region of the accompanying ^1^H-^15^N CP spectrum (Figure S2a) is featureless − due to limited chemical shift dispersion, this region is particularly prone to loss of resolution as a result of conformational heterogeneity. Together, the CP spectra of isolated N-FAT10-WT suggest a loss of structural organization upon dense packing.

**Figure 1.**
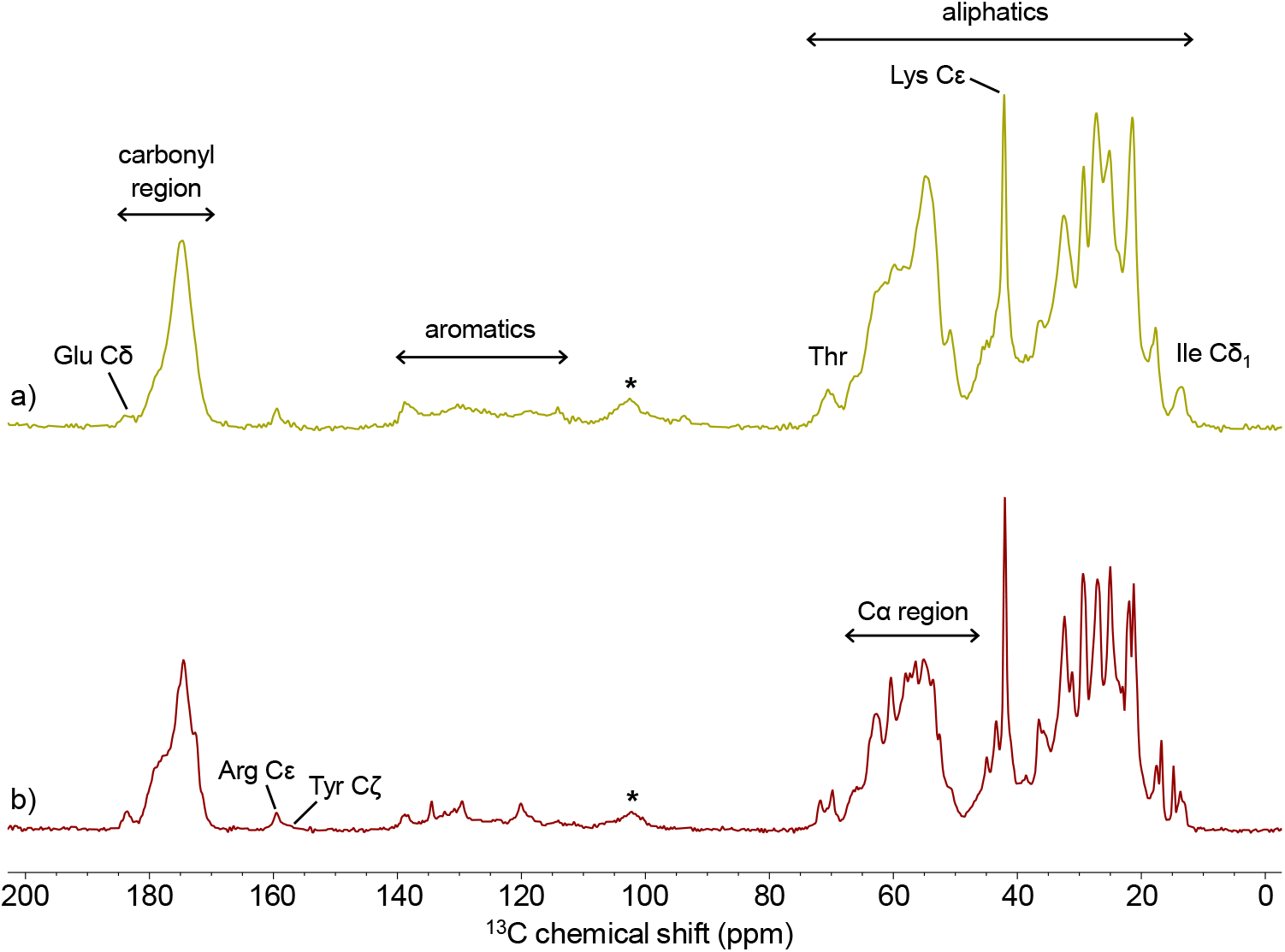
^1^H-^13^C cross-polarization spectra of (a) N-FAT10-WT and (b) N-FAT10-WT co-sedimented with NUB1L. Spectra were recorded at 18.8 T with a spinning frequency of 14.5 kHz. The sample temperature was 4 °C. The stars indicate spinning sidebands.

The ^13^C-^13^C correlation spectrum of isolated N-FAT10-WT, recorded with a prolonged mixing time of 50 ms, is shown in Figure 2a. Broad cross peaks are observed from at least Ala, Ile, Leu, Lys, Pro, Ser, Thr, and Val. For none of these residues, distinct chemical environments can be resolved (Figure 2b). To guide the eye, we marked the average chemical shifts of these residues, depending on whether the local secondary structure is a helix, coil, or sheet. For this, we used the values provided in the supplementary table of Fritzsching *et al*.^61^, who extracted them from the PACSY database^62^. The chemical shifts of the C_α_ atoms experience the strongest perturbations, since they are part of the backbone. The average chemical shifts of the coil and sheet conformations tend to be similar, while the helical conformation stands out. We observe that the broad cross peaks in Figure 2a are mainly located in coil/sheet regions, suggesting that these are the dominant conformations in the sample. Indeed, increased β-sheet content is a characteristic of protein aggregates.^63^ The ^13^C-^13^C correlation spectrum is thus in agreement with the formation of an amorphous protein aggregate upon dense packing of isolated N-FAT10-WT.

**Figure 2.**
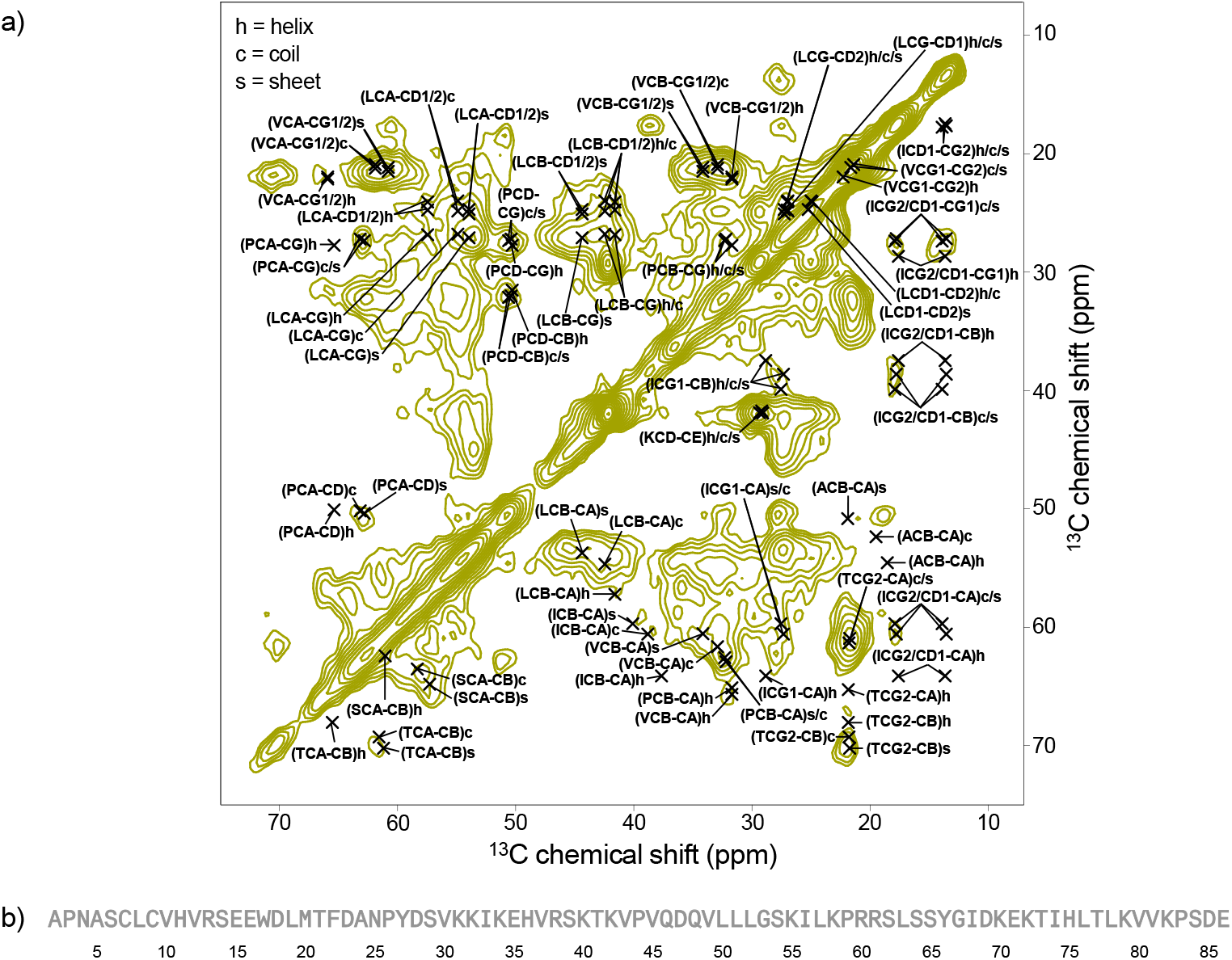
(a) ^13^C-^13^C DARR spectrum of isolated, densely packed N-FAT10-WT, recorded at 18.8 T. Average chemical shifts of Ala, Ile, Leu, Lys, Pro, Ser, Thr, and Val in helix (h), coil (c), or sheet (s) conformation are indicated. Below the diagonal, C_α_-C_β_ cross peaks, cross peaks of Ile and Thr, and the C_δ_-C_ε_ cross peak of Lys are shown. Above the diagonal, the other cross peaks of Leu, Pro, and Val are shown. (b) Sequence of N-FAT10-WT (amino acids 2-86). Sequential assignment was not possible for any of the residues.

The ^1^H-^13^C CP spectrum of N-FAT10-WT co-sedimented with NUB1L is plotted in Figures 1b (bordeaux). The spectrum now shows well-defined resonances, particularly from aromatic and aliphatic sidechains. Resolved signals from Thr and Ile are visible at the edges of the aliphatic region. The carbonyl and C_α_ regions, however, are not as nicely resolved as for isolated N-FAT10-C0 in the β-grasp fold (see Ref. ^44^). The ^1^H-^15^N CP spectrum (Figure S2b) shows a similar phenomenon: a set of well-defined resonances, including from Pro and the sidechain of His, combined with (or on top of) a smooth backbone region. Both CP spectra suggest that if N-FAT10-WT is co-sedimented with NUB1L, it adopts, in part, a regular, stable structure. This is confirmed by ^13^C-^13^C and ^15^N-^13^C correlation spectra of the same sample (Figures 3a,b). Well-resolved cross peaks are observed, but their number is too small to account for all 82 residues present.

**Figure 3.**
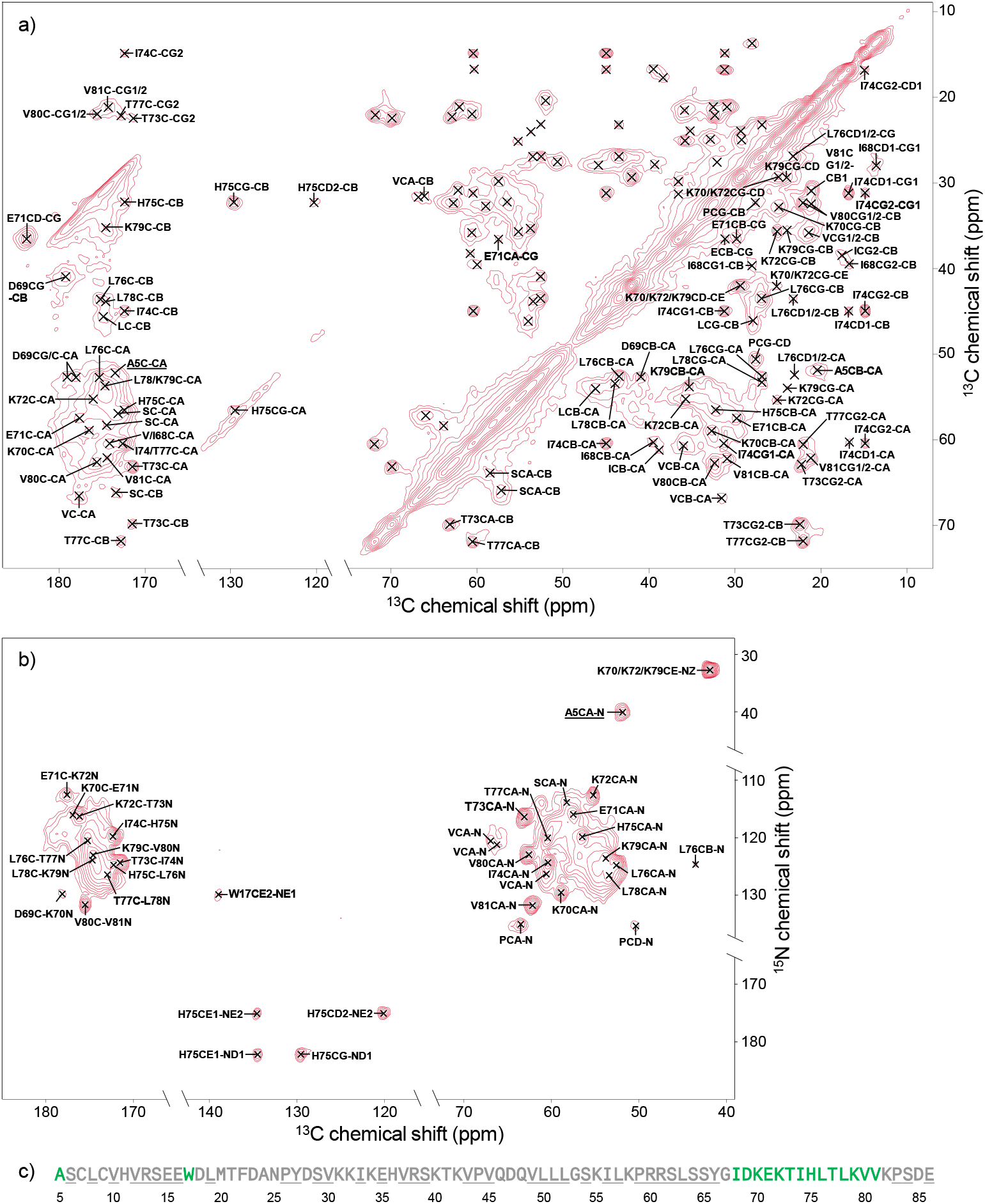
(a)^13^C-^13^C DARR spectrum and (b) ^15^N-^13^C ZF TEDOR spectrum of N-FAT10-WT co-sedimented with NUB1L. Both spectra were recorded at 18.8 T. Assigned cross peaks are marked and labeled. For the strip plot demonstrating the sequential assignments, see Figure S4. (c) Overview of sequential assignments of N-FAT10-WT co-sedimented with NUB1L. Green = unambiguous and grey = unobserved. Of the grey, underlined residues, we observe 1xE, 1xI, 1xL, 1xP, 1xR, 2xS, 3xV, 1xY.

The ^13^C-^13^C and ^15^N-^13^C correlation spectra in Figures 3a,b provide informative fingerprints but do not allow sequential assignment of the residues of U-^13^C,^15^N-N-FAT10-WT. For this purpose, ^15^N-^13^C-^13^C correlation spectra are necessary. Given the relatively low yield of N-FAT10-WT and the fact that, upon co-sedimentation, NUB1L takes up most of the space in the MAS rotor, recording these spectra is, with a standard MAS probe, rather time consuming. We therefore decided to explore the benefits of the MAS CryoProbe. First, to verify reproducibility and the integrity of a newly co-sedimented sample, we acquired ^15^N-^13^C NCO and NCA spectra, see Figure S3. Agreement with the ZF TEDOR spectrum in Figure 3b is very good. Any differences are traced back to the use of distinct pulse sequences and acquisition parameters: (1) In the NCO and NCA experiments, magnetization is transferred specifically from nitrogen nuclei to C’ and C_α_ nuclei in the backbone. Hence, no cross peaks are observed from the sidechains of His, Arg, and Lys. (2) In the NCO and NCA experiments, magnetization transfer is slightly more efficient, as illustrated by the cross peaks with the C_β_ nuclei of T73, I74, H75, and T77 in Figure S3. (3) Cross peaks with the C_α_ and C_δ_ nuclei of Pro are evident at (63.5,135.2) and (50.6,135.2) ppm in the ZF TEDOR spectrum of Figure 3b, but absent in the NCO and NCA spectra of Figure S3. The reason is the inefficient ^1^H-^15^N cross-polarization to the nitrogen of Pro at the start of the NCO and NCA sequences, which is circumvented in ZF TEDOR because the sequence starts with a ^1^H-^13^C cross-polarization step^50^.

Next, ^15^N-^13^C-^13^C correlations were obtained in separate NCOCX and NCACX experiments. Sequential assignment was possible for amino acid residues I68 through V81; the strip plot is provided in Figure S4. Cross peaks are mostly limited to C’, C_α_, and C_β_ nuclei and patterns of individual residues in the ^13^C-^13^C and ^15^N-^13^C correlation spectra were very helpful to confirm site-specific assignments. In addition to I68-V81, a series of residues with missing backbone connectivity is observed: 1xA, 1xE, 1xI, 1xL, 1xP, 1xR, 2xS, 3xV, 1xY, and 1xW. Since there is only one Trp in the sequence, the last of these must be W17. The unusual ^15^N chemical shift of the Ala cross peak, at (52.0,40.1) ppm in Figure 3b, pins it down as the N-terminus (A5).^64^ The C’-C_α_ cross peak of A5 is also visible in Figure 3a, at (173.6,52.0) ppm. An overview of the assignments of N-FAT10-WT co-sedimented with NUB1L is provided in Figure 3c. Chemical shift values are provided in Table S3.

The cross peaks from the N-termini constitute the only significant difference between the MAS NMR spectra presented here for N-FAT10-WT co-sedimented with NUB1L and the spectra reported previously^44^ for stabilized N-FAT10-C0 co-sedimented with NUB1L. As the result of the use of TEV protease for cleavage of the purification tag, the N-terminal residue of N-FAT10-C0 is a Gly, not an Ala and the chemical shifts of the associated cross peaks differ accordingly. Otherwise, the complexes of N-FAT10-WT and N-FAT10-C0 of FAT10 with NUB1L appear identical. For completeness, we provide the TALOS-N predictions of the torsion angles and secondary structures for the N-FAT10-WT complex in Figure S5 and the structure prediction by AlphaFold-Multimer in Figure 4. There are no significant changes. The inset in Figure 4 shows a projection of the empirical secondary structure predictions onto the AlphaFold-Multimer prediction. As for the N-FAT10-C0 complex, chemical shifts indicate that the intermolecular β-sheet is, in reality, three residues longer: residues T73-V81 are classified as strand whereas AlphaFold-Multimer suggests that only residues L76-V81 of N-FAT10-WT are involved. Residues I68-K72 form a loop.

**Figure 4.**
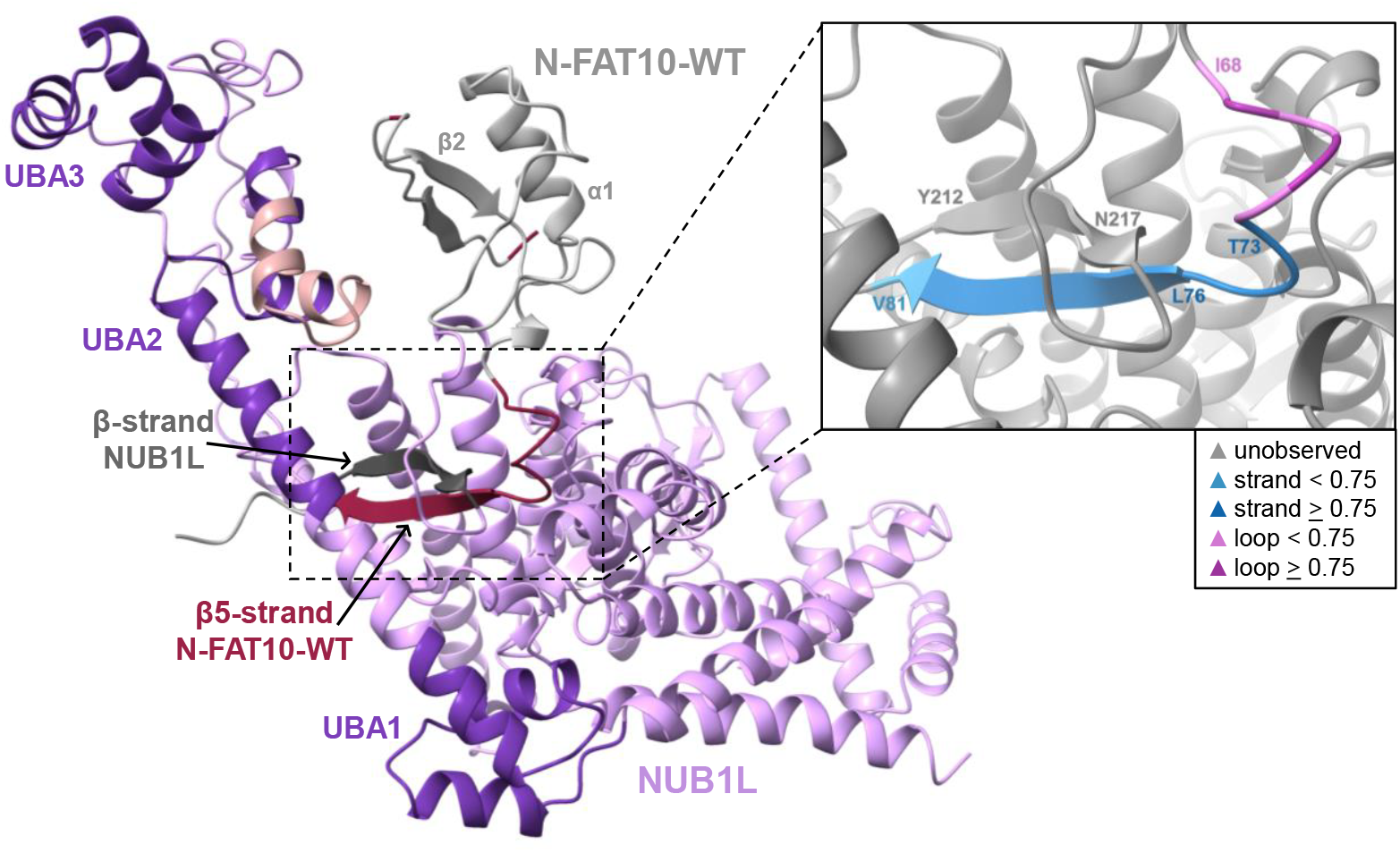
Best-ranked AlphaFold-Multimer structure prediction (confidence score of 0.77) for the interaction of N-FAT10-WT (amino acids 5-86) and NUB1L. The residues of N-FAT10-WT that are observed by MAS NMR are highlighted in red. The inset shows the empirical secondary structure prediction for residues I68-V81 projected on the AlphaFold-Multimer prediction (see Figure S5 for the backbone torsion angles and details). MAS NMR confirms the existence of the intermolecular β-sheet and shows that it is, in reality, three residues longer. Integrity of the other secondary structure elements in N-FAT10-WT is negated by MAS NMR. The residues of NUB1L highlighted in rosa are absent in the shorter splice variant NUB1.

## DISCUSSION

In solution, FAT10 exists in both folded and unfolded states.^39,40,42^ The insight that NUB1(L) binds FAT10 in a mostly unfolded state in preparation of rapid degradation by the proteasome^42,44^ reveals the biological function of this property, at least for the N-domain. Hydrophobic residues that are exposed while sampling an unfolded state readily interact with exposed hydrophobic residues of other proteins or FAT10 copies, causing FAT10 to be prone to aggregation. This may not be just a detrimental side effect: FAT10 seems able to reduce the thermodynamic stability of its substrates, with the effect of speeding up their degradation.^39^ In the presence of NUB1(L), complex formation reduces the chance that two unfolded FAT10 molecules interact with each other and aggregation is expected to be mitigated. In our handling of both proteins, we have observed that NUB1L can recover N-FAT10-WT that has already started to aggregate: upon addition of NUB1L, white, cloudy streaks disappear and the solution becomes clear again.

MAS NMR experiments indicate that the structures of the complexes of N-FAT10-WT and N-FAT10-C0 with NUB1L are the same. This is not an obvious, predictable outcome. The removal of the Cys residues effectively stabilizes the N-domain of FAT10 in the ubiquitin-like β-grasp fold, as evidenced by the appearance of residues P59-T73 in the HSQC spectrum (G53 and S54 are absent instead) and the possibility to form crystals.^31^ Also, Arkinson *et al*. could not detect complex formation between full-length FAT10-C0 and NUB1,^42^ which explains why removal of the Cys residues hampers degradation^31^. Possibly binding is abrogated because, in the presence of the (stabilized) C-domain, bulky, folded parts, on both ends of the β-sheet-forming stretch (T73-V81) prevent insertion into the looped-out portion of NUB1(L). The shortening of the second ubiquitin-associated (UBA2) domain in NUB1 (Figure 4) might play a role as well. With dissociation constants of 176 nM^42^ and 813 nM^65^, respectively, full-length wild-type FAT10 binds more tightly to NUB1 than to NUB1L, perhaps at the expense of flexibility. Arkinson *et al*. have argued, based on the similarity of the characteristic times of folding/unfolding and degradation, that NUB1 uses conformational selection to bind and trap the spontaneously unfolded FAT10. Our observation that stabilized N-FAT10-C0 forms the same complex with NUB1L as N-FAT10-WT, could argue for the opposite, namely binding-induced unfolding. This would be in line with the observation that in isolated N-FAT10-C0 the residues I68-L76 are primed for capture of the β5-strand^44^.

The equivalence of the wild-type and Cys-free complexes validates the biological relevance of the previous MAS NMR investigations of N-FAT10-C0^44^. It also lends further support for the fuzzy-complex model. We previously proposed that the N-terminus, W17, and the eleven more residues that, due to missing backbone connectivity in the three-dimensional spectra, could not be assigned sequentially, are anchor points where N-FAT10-C0 interacts non-covalently with NUB1L in between disordered stretches of residues. Here, in the wild-type complex, we observe these residues exactly as before, proving that they are not an artifact of incomplete interaction with stabilized N-FAT10-C0. Importantly, we confirm that the N-terminus of N-FAT10, regardless of whether it is a Gly or an Ala, takes on a regular, stable structure in the complex with NUB1L. We do not yet know where on NUB1(L) the N-terminus binds, but we nevertheless suspect that this binding is a critical aspect of how FAT10 and NUB1L together enable the rapid degradation of substrates. The N-terminus is the first residue of the unfolded FAT10 chain to enter the proteasome^23,31,42^ and, hence, control over its position (and not over the far-away β5-strand) seems essential to initiate degradation.

Ultracentrifugation has long been used in biomolecular MAS NMR to densely pack insoluble systems like protein microcrystals, amyloid fibrils, viral capsids, and membrane proteins in the MAS rotor. The realization that ultracentrifugation, first *ex situ* in an ultracentrifuge from solution into the MAS rotor and second *in situ* in the spectrometer by MAS, can also be used to generate a dense phase of soluble proteins is more recent.^66–68^ High viscosity and sedimentation work together to immobilize proteins and prevent anisotropic interactions from averaging out, at least on the NMR time scale.^45,69^ Initially, the method was applied to large (∼500 kDa) and (to keep spectral complexity under control) multimeric protein systems because sedimentation theory predicts that large systems are immobilized most effectively. The use of co-sedimentation to form large molecular complexes has also been demonstrated.^46,70^ Mainz *et al*. succeeded in co-sedimenting the chaperone αB-crystallin with lysozyme, which was first unfolded with an excess of TCEP,^71^ a known trick to aid the formation of chaperone-client complexes^72^. Over the years, it has become clear that sedimentation by ultracentrifugation also works well for much smaller proteins^73^. This has recently been highlighted by Bell *et al*., who reported MAS NMR spectra of soluble proteins in the size range 40-140 kDa.^74^ While solution NMR of proteins of this size is certainly possible, it requires tailored isotope labelling and dedicated pulse sequences. MAS NMR provides a straightforward and informative alternative, especially with the enhanced sensitivity of the MAS CryoProbe.

Here, we have shown that simple mixing in solution followed by ultracentrifugation is a viable method for the preparation of small protein complexes in the dense phase; the combined molecular weight of the N-FAT10-WT/NUB1L complex is 80.3 kDa (70.4 kDa from natural abundance NUB1L and 9.9 kDa from U-^13^C,^15^N-N-FAT10-WT). Because unbound N-FAT10-WT is prone to aggregation, we took care to keep its concentration low during the procedure, at 1 mg/mL. The co-sediment has a yellowish, semi-transparent appearance and a sticky texture, which makes it somewhat difficult to handle (see the footnote at Table S1). However, once the co-sediment has undergone the strong ultracentrifugation of MAS, it is stable for many months, as verified by MAS NMR (storage in between experiments was at 4 °C) and in agreement with reports by others^75^.

For the purpose of co-sedimentation, we prepared a co-solution of equal starting concentrations (in mol/L) of N-FAT10-WT and NUB1L, with the aim of producing the maximum concentration of the complex relative to the concentration of the two unbound proteins. According to basic sedimentation theory, larger, heavier proteins sediment faster during ultracentrifugation (or MAS), which means that a complex is extracted from the solution before unbound proteins.^45^ As a consequence, the equilibrium in the solution will shift and, in this way, co-sedimentation could foster complex formation. It would be interesting to investigate to what extent this mechanism has contributed here and if it could aid the formation of, for example, chaperone-client complexes^13^. The dissociation constant for N-FAT10-WT and NUB1L (813 nM) is on the low end of what has been observed for chaperone-client interactions^76^ and the kinetics are with *k*_on_ = 1600 M^-1^s^-1^ relatively slow^42^. Both these properties are probably beneficial, the latter because a complex that stays intact longer is more easily dragged into the sedimented layer where it cannot readily dissociate.

## CONCLUSION

MAS NMR of N-FAT10-C0 and, as reported above, of N-FAT10-WT co-sedimented with NUB1L has provided critical insight into their interaction. The formation of an intermolecular β-sheet has been experimentally verified and the AlphaFold-Multimer structure prediction has been amended. NUB1L acts as a holdase for N-FAT10-C0/WT: it holds N-FAT10-C0/WT in a conditional, but incomplete fold. Further investigation by MAS NMR of an alternatively labeled sample should provide information about binding locations on NUB1L and its structure. When we started this project, the use of MAS NMR was a leap into the unknown but it has turned out well. Co-crystallization of N-FAT10-C0/WT and NUB1L would probably have been difficult because such a large part of N-FAT10-C0/WT is disordered. Cryo-electron microscopy would be expected to struggle with the small size and, when bound to the proteasome^42^, the complex seems too dynamic for high-resolution images. Moreover, neither of these two methods can probe local chemical environments or dynamics, but with MAS NMR these are easily accessible. Hence, we advocate MAS NMR for the investigation of fuzzy complexes like those formed between chaperone and client and, more generally, of conditional folds of IDPs and IDRs.

## Associated content

### Supporting Information

Protocol for expression and purification of U-^13^C,^15^N-N-FAT10-WT including SDS-PAGE analysis; overview of MAS NMR experiments and acquisition parameters; ^1^H-^15^N cross-polarization spectra; ^15^N-^13^C NCO and NCA spectra acquired with the MAS CryoProbe; representative strip plot of ^15^N-^13^C-^13^C NCOCX and NCACX spectra; table with ^15^N and ^13^C chemical shifts of N-FAT10-WT in complex NUB1L, torsion angle and secondary structure predictions.

### Data availability

All data generated in this study have been deposited in the [Zenodo repository](https://doi.org/10.5281/zenodo.18290249). The NMR assignments for N-FAT10-WT in complex with NUB1L have been deposited to the Biological Magnetic Resonance Data Bank (BMRB) under accession code 53486.

## Acknowledgements

The authors are deeply indebted to Prof. Dr. Marcus Groettrup. Without his enthusiasm and scientific excellence, this work would not have been possible. He is sorely missed. This research was funded by the Deutsche Forschungsgemeinschaft through the Emmy Noether Program (Project No. 321027114), SFB 969 (Project No. 189682160), and SFB 1527 (Project No. 454252029). The authors thank Michael Kovermann, Ulrich Haunz, and Anke Friemel for maintenance of the NMR instrumentation, Gunter Schmidtke for support in the biolab, and Stefan Gerlach for support regarding the Scientific Compute Cluster at the University of Konstanz (SCCKN).

## Notes

### Competing Interest Statement

The authors have declared no competing interest.

### Summary of Updates

Figures 2, 3, 4, S3, S4, and S5 have been updated

