## Supporting Information for "Co-sedimentation is the key to the structural investigation of wild-type FAT10"

### 1. Expression and purification of U-$^{13}C$,$^{15}N$-N-FAT10-WT

Molecular cloning was achieved by the classical restriction enzyme approach using BsaI and XhoI. Wild-type N-domain of human FAT10 (amino acids 2-86) and an N-terminally truncated variant thereof (amino acids 5-86) were cloned into pSUMO vectors for expression as $His_6$-SUMO-fusion proteins. Isolated candidate clones were verified by DNA sequencing (Microsynth). Expression was performed in *E. coli* BL21-CodonPlus(DE3)-RIPL competent cells (Agilent Technologies). For uniform $^{13}C$ and $^{15}N$ labelling, 3.6 g of U-$^{13}C_6$-D-glucose and 0.5 g of $^{15}N$-ammonium chloride were added per litre of M9 minimal medium as exclusive sources of carbon and nitrogen. Bacteria cells were grown at 37 °C to an $OD_{600}$ of 0.5-0.6, induced with 0.4 mM IPTG at 21 °C overnight and harvested by centrifugation (RCF 4000 g, 10 min, 8 °C). Harvested cells were lysed in lysis buffer (20 mM TRIS-HCl (pH 8.0), 300 mM NaCl, 10 mM imidazole, 1 mM TCEP, 10 % v/v glycerol, 0.1 % v/v Triton X-100, 100 µg/mL lysozyme, 1 mM PMSF and EDTA-free protease inhibitor). After sonication, cell debris was removed by centrifugation (RCF 47000 g, 30 min, 8 °C). The supernatant was filtered and applied to Ni-NTA Agarose beads (Macherey-Nagel) for affinity chromatography. The protein was eluted with elution buffer (20 mM TRIS-HCl (pH 8.0), 300 mM NaCl, 500 mM imidazole and 1 mM TCEP) and buffer exchanged to binding buffer (20 mM TRIS-HCl (pH 8.0), 300 mM NaCl, 10 mM imidazole and 1 mM TCEP). His-tagged Ulp1 protease was allowed to act overnight (4 µg of protease per 1 mg of recombinant protein). The Ulp1 protease recognizes the three-dimensional fold of the SUMO-tag preserving the sequence of N-FAT10, which starts, for both variants, with an alanine residue. After cleavage, His-tagged Ulp1 protease and cleavage byproducts were separated by a second Ni-affinity chromatography using binding buffer. The volume of the flow-through containing U-$^{13}C$,$^{15}N$-N-FAT10-WT (amino acids 2-86 or 5-86) was reduced to 5 mL and filtered. Final purification (Figure S1) was achieved by size-exclusion chromatography (Cytiva, formerly Amersham Biosciences, ÄKTA pure chromatography system equipped with a HiLoad 16/600 Superdex 75 pg column) yielding up to 3 mg per litre of M9 medium. Before preparation of the samples for MAS NMR, U-$^{13}C$,$^{15}N$-N-FAT10-WT (amino acids 2-86 or 5-86) was in 20 mM HEPES (pH 7.5), 150 mM NaCl and 1 mM TCEP and concentrated to specified values (Amicon Ultra-15 centrifugal filter unit, MWCO 3 kDa).

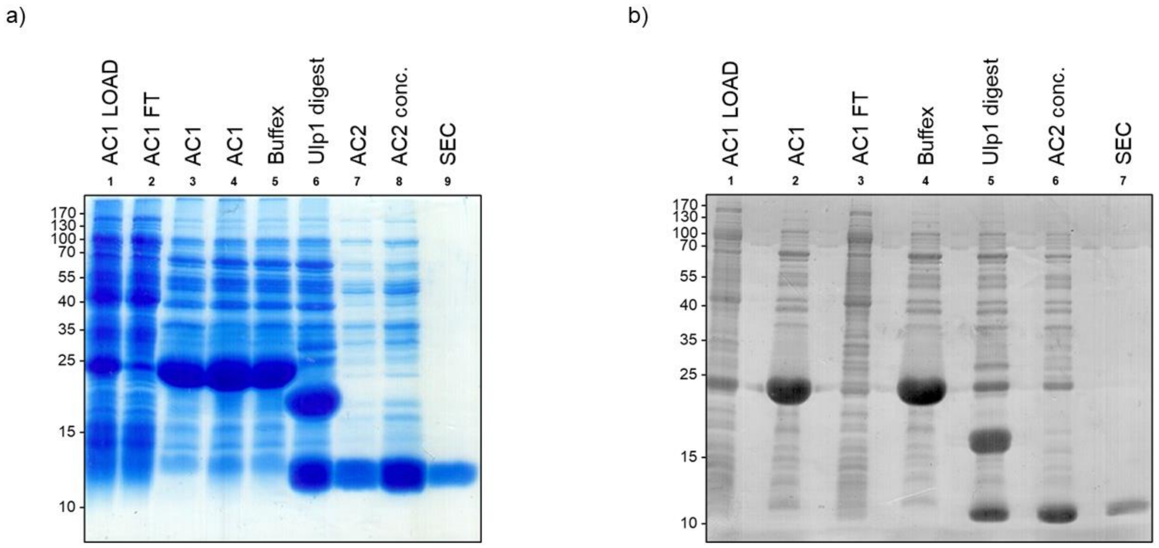

***Figure S1:*** *SDS-PAGE analysis for the purifications of (a) N-FAT10-WT, amino acids 2-86 and (b) N-FAT10-WT, amino acids 5-86. 5 µL of gel sample buffer with SDS (reducing conditions) were added to 20 µL of each sample. After boiling for 5 min at 95 °C, samples were loaded on a 15 % polyacrylamide gel and separated applying for 18-20 min a constant voltage of 120 V and for 55-65 min a constant voltage of 160 V followed by colloidal Coomassie staining. For molecular weight estimation, PageRuler Prestained Protein Ladder size standards (Thermo Fisher) were used. AC = Affinity Chromatography, FT = Flow-Through, Buffex = Buffer exchange, SEC = Size-Exclusion Chromatography.*

#### 2. MAS NMR spectroscopy

A first series of experiments was performed at 18.8 T ($^{1}$H Larmor frequency of 800.3 MHz) in the NMR Core Facility at the University of Konstanz. The console was a Bruker Avance NEO. A Bruker 3.2 mm E-free HCN MAS probe was used, together with regular-wall 3.2 mm zirconia rotors (maximum sample volume 32 µL). A second series of experiments was performed at 14.1 T ($^{1}$H Larmor frequency of 600.3 MHz) at the site of Bruker Switzerland in Fällanden. The console was again a Bruker Avance NEO. A Bruker MAS CryoProbe was used, together with a dedicated thin-wall 3.2 mm silicon nitride rotor (maximum sample volume 87 µL, for our measurements reduced to 47 µL by placing a 5-mm spacer at the top and a 2-mm spacer at the bottom of the rotor). Table S1 provides an overview of all experiments on both instruments. Calibration of the Bruker Cooling Unit (BCU) II was performed by monitoring the chemical shift of $^{79}$Br in KBr powder.[1] The $^{13}$C chemical shifts are referenced indirectly to DSS in $D_2O$ (0.5 % by weight),[2] i.e., the $^{13}$C adamantane methylene peak is observed at 40.49 ppm. The $^{15}$N chemical shifts are referenced to liquid ammonia at 25 °C.[3]

***Table S1:*** *MAS NMR experiments per sample.*

| Experiment | Field (T) | Number of scans | Non-uniform sampling | $^{15}N$-$^{13}C$ or $^{13}C$-$^{13}C$ mixing time (ms) (scheme) | Temperature (°C) | MAS frequency (kHz) | Measurement time |
|---|---|---|---|---|---|---|---|
| **U-$^{13}C$,$^{15}N$-N-FAT10-WT (2-86)** | | | | | | | |
| hC | 18.8 | 512 | | | 4 | 14.5 | |
| hN | 18.8 | 256 | | | 4 | 14.5 | |
| hCC | 18.8 | 64 | | 50 (DARR) | 4 | 14.5 | 15.5 h |
| **U-$^{13}C$,$^{15}N$-N-FAT10-WT (5-86) in complex with natural abundance NUB1L** | | | | | | | |
| hC | 18.8 | 256 | | | 4 | 14.5 | |
| hN | 18.8 | 512 | | | 4 | 14.5 | |
| hCC | 18.8 | 128 | | 10 (DARR) | 4 | 14.5 | 3 d 0.5 h |
| hNC | 18.8 | 512 | | 1.1 (ZF TEDOR) | 4 | 14.5 | 3 d 8 h |
| **U-$^{13}C$,$^{15}N$-N-FAT10-WT (5-86) in complex with natural abundance NUB1L (MAS CryoProbe rotor)#** | | | | | | | |
| NCO | 14.1 | 128 (2x) | | | 4 | 12.0 | 3.5 h (2x) |
| NCA | 14.1 | 128 (2x) | | | 4 | 12.0 | 3.5 h (2x) |
| NCOCX | 14.1 | 256 (2x) | 42 % | 50 (CORD) | 4 | 12.0 | 3 d 4 h (2x) |
| NCACX | 14.1 | 384/376 | 42 % | 50 (CORD) | 4 | 12.0 | 4 d 18.5 h/ 4 d 16 h |

*#Co-sedimented N-FAT10-WT and NUB1L was transferred from an ultracentrifuge tube to the MAS CryoProbe rotor using a benchtop centrifuge. This transfer was not without incident due to the sticky texture of the pellet and an unscheduled delay between ultracentrifugation and transfer, which caused an uptake of buffer. We estimate that the amount of U-$^{13}C$,$^{15}N$-N-FAT10-WT in the MAS CryoProbe rotor was 6-9 times less than what we had in the standard MAS rotor, i.e., less than 0.5 mg. Still, with the additional help of non-uniform sampling, it was possible to record the NCOCX and NCACX spectra required for sequential assignment.*

Typical $\pi/2$-pulse lengths were 2.8-3.0 μs for $^{1}H$, 4.1-5.0 μs for $^{13}C$ and 6.0-9.0 μs for $^{15}N$. $^{1}H$-$^{13}C$ and $^{1}H$-$^{15}N$ cross-polarization (CP)[4] are realized using a linearly ramped radio frequency (RF) field on the $^{1}H$ channel from 90 to 100 % amplitude; the centre of the ramp was set to match the n = +1 Hartmann-Hahn condition.[5] Swept-frequency two-pulse phase modulation ($SW_f$-TPPM)[6] or small phase incremental alternation with 64 steps (SPINAL-64)[7] were applied for proton decoupling. The States-TPPI method[8] was used for phase-sensitive detection in all indirect dimensions. During $^{13}C$-$^{13}C$ DARR mixing and the z-filter delay in $^{15}N$-$^{13}C$ ZF TEDOR, protons were irradiated at an RF field matching the spinning frequency of 14.5 kHz. The z-filter delays were set to ~200 μs, matching a multiple of the rotor period. Table S2 shows the acquisition parameters per experiment and sample. To record the NCOCX and NCACX spectra, exponentially biased random sampling with a preference for short t1- and t2-values was used. Two experiments were performed with sparse sampling of 25 %, one with a NUS-$T_2$ of 1 s and one with a NUS-$T_2$ of 3 ms. Adding the two experiments gives a combined sparse sampling of 42 %.

For free induction decays with unnecessary long acquisition times, the number of effective points was adjusted accordingly. 1D spectra were zero-filled and baseline corrected, but no apodization was applied. For 2D and 3D spectra, 60°- or 72°-shifted squared-sine bell apodization (corresponding to an offset of 0.33 or 0.40 in NMRPipe) and zero filling were used in all dimensions.

***Table S2:*** *Acquisition parameters for MAS NMR experiments.*

| Experiment | CP contact time (ms) | | Radio frequency field strength (kHz) | | | | | | | | Dwell time (μs) | | | Acquisition time (ms) | | | Carrier position (ppm) | | | Recycle delay (s) |
|---|---|---|---|---|---|---|---|---|---|---|---|---|---|---|---|---|---|---|---|---|
| | $^{1}$H-$^{13}$C/ $^{1}$H-$^{15}$N CP | $^{15}$N-$^{13}$C DCP | $^{1}$H-$^{13}$C/$^{1}$H-$^{15}$N CP | | $^{15}$N-$^{13}$C DCP | | $\pi$/2- and $\pi$-pulses | | | Decoupling | | | | | | | | | | |
| | | | $^{1}$H (linear)# | $^{13}$C/$^{15}$N | $^{15}$N | $^{13}$C (tangent)# | $^{1}$H | $^{13}$C | $^{15}$N | $^{1}$H | $^{13}$C | $^{13}$C | $^{15}$N | $^{13}$C | $^{13}$C | $^{15}$N | $^{13}$C | $^{13}$C | $^{15}$N | |
| **U-$^{13}$C,$^{15}$N-N-FAT10-WT (2-86)** | | | | | | | | | | | | | | | | | | | | |
| hC | 1.2 | | 71.1 | 50.0 | | | 83.3 | | | 71.4, SPINAL-64 | 7.00 | | | 14.3 | | | 100 | | | 2.5 |
| hN | 1.0 | | 67.4 | 50.0 | | | 83.3 | | | 71.4, SPINAL-64 | | | 12.0 | | | 12.3 | | | 104 | 3.5 |
| hCC | 1.2 | | 71.1 | 50.0 | | | 83.3 | 50.0 | | 71.4, SPINAL-64 | 11.6 | 23.0 | | 11.9 | 4.8 | | 100 | 100 | | 2.0 |
| **U-$^{13}$C,$^{15}$N-N-FAT10-WT (5-86) in complex with natural abundance NUB1L** | | | | | | | | | | | | | | | | | | | | |
| hC | 1.0 | | 67.8 | 50.0 | | | 83.3 | | | 71.4, SPINAL-64 | 6.1 | | | 25.0 | | | 100 | | | 3.0 |
| hN | 0.9 | | 59.1 | 41.7 | | | 83.3 | | | 71.4, SPINAL-64 | | | 12.0 | | | 24.6 | | | 104 | 3.0 |
| hCC | 1.0 | | 67.8 | 50.0 | | | 83.3 | 50.0 | | 71.4, SPINAL-64 | 11.3 | 23.0 | | 23.1 | 11.4 | | 100 | 100 | | 2.0 |
| hNC | 1.0 | | 67.8 | 50.0 | | | 83.3 | 50.0 | 27.8 | 83.3/71.4, SW$_f$-TPPM | 12.0 | | 69.0 | 24.6 | | 7.6 | 100 | | 104 | 2.5 |
| **U-$^{13}$C,$^{15}$N-N-FAT10-WT (5-86) in complex with natural abundance NUB1L (MAS CryoProbe rotor)** | | | | | | | | | | | | | | | | | | | | |
| NCO | 0.75 | 1.25 | 68.2 (tangent) | 41.7 | 6.95 | 6.05 | 85.9 | | | 90.0, SW$_f$-TPPM/ CW | 11.0 | | 333.3 | 20.0 | | 8.0 | 110† | | 122 | 2.0 |
| NCA | 0.75 | 1.75 | 68.8 (tangent) | 41.7 | 3.66 | 20.8 | 85.9 | | | 90.0, SW$_f$-TPPM/ CW | 11.0 | | 333.3 | 20.0 | | 8.0 | 110† | | 122 | 2.0 |
| NCOCX | 0.75 | 1.25 | 68.2 (tangent) | 41.7 | 6.95 | 6.05 | 85.9 | 60.6 | 41.7 | 90.0, SW$_f$-TPPM/ CW | 9.5 | 333.3 | 333.3 | 19.5 | 8.0 | 8.0 | 110 | 110† | 122 | 1.8 |
| NCACX | 0.75 | 2.0 | 72.3 (tangent) | 41.7 | 3.66 | 21.4 | 89.3 | 60.6 | 41.7 | 95.0, SW$_f$-TPPM/ CW | 9.5 | 333.3 | 333.3 | 19.5 | 8.0 | 8.0 | 110 | 110† | 122 | 1.8 |

*#Corresponds to 100 % of the ramp. †For $^{15}$N-$^{13}$C SPECIFIC-CP, the CO carrier was centred at 172 and 174 ppm for NCO and NCOCX experiments, respectively. The CA carrier was centred at 55 and 56 ppm for NCA and NCACX experiments, respectively.*

#### 3. $^{1}H$-$^{15}N$ cross-polarization spectra

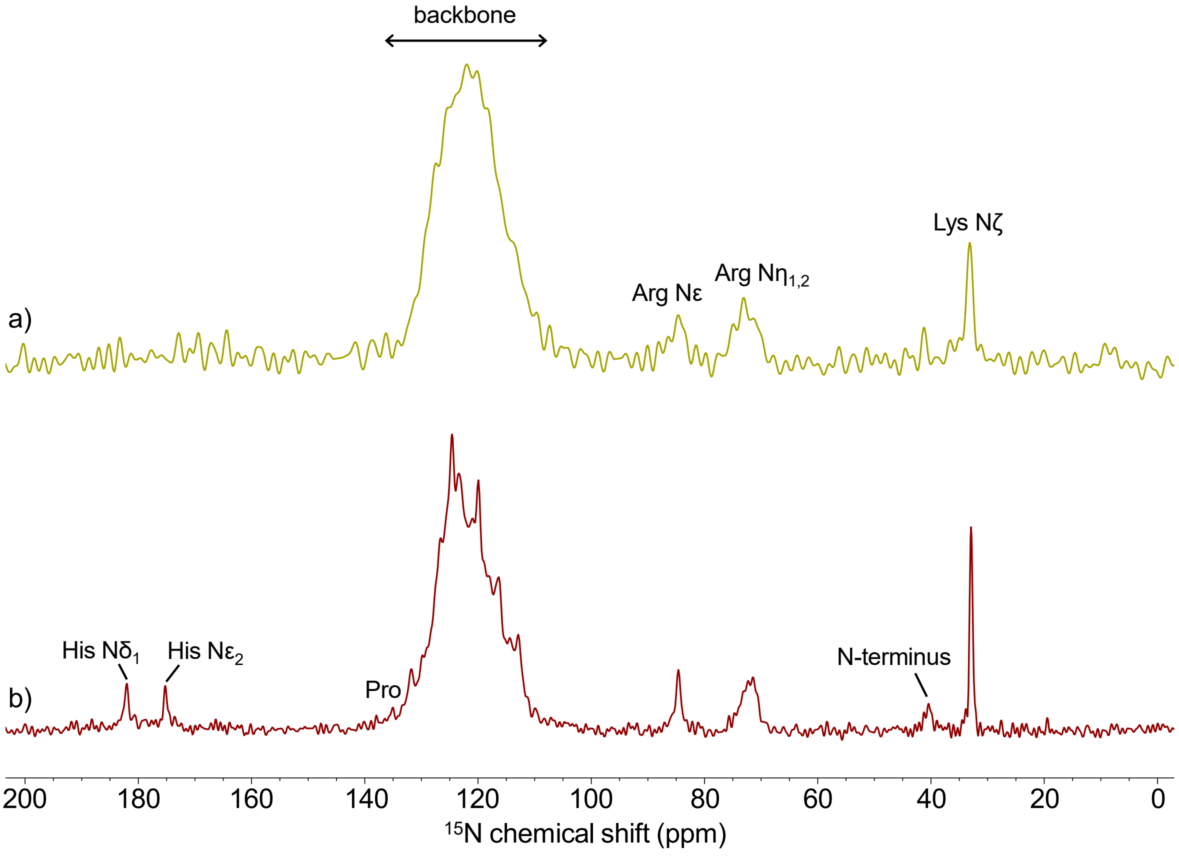

***Figure S2:*** *$^{1}H$-$^{15}N$ cross-polarization spectra of (a) N-FAT10-WT and (b) N-FAT10-WT co-sedimented with NUB1L. Spectra were recorded at 18.8 T with a spinning frequency of 14.5 kHz.*

#### 4. $^{15}N$-$^{13}C$ correlation spectra of N-FAT10-WT in complex with NUB1L

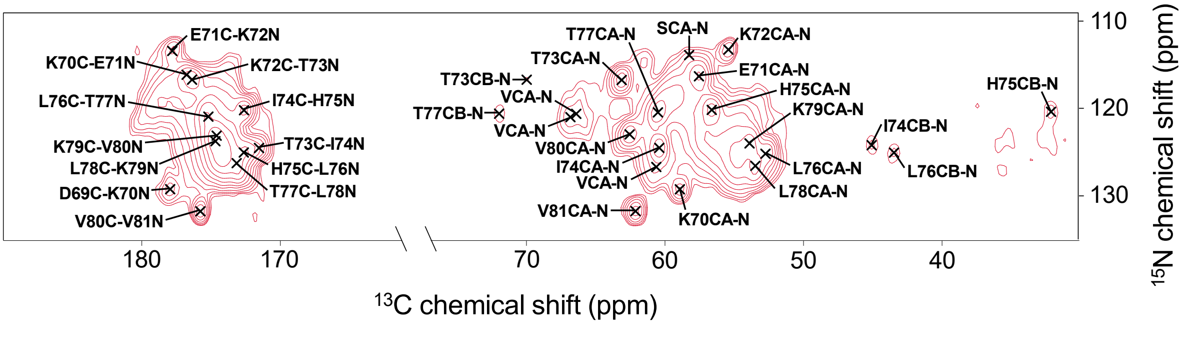

***Figure S3:*** *$^{15}N$-$^{13}C$ NCO and NCA spectra of N-FAT10-WT co-sedimented with NUB1L. Spectra were recorded at 14.1 T with a spinning frequency of 12.0 kHz. Assigned cross peaks are marked and labeled.*

#### 5. Strip plot of N-FAT10-WT in complex with NUB1L

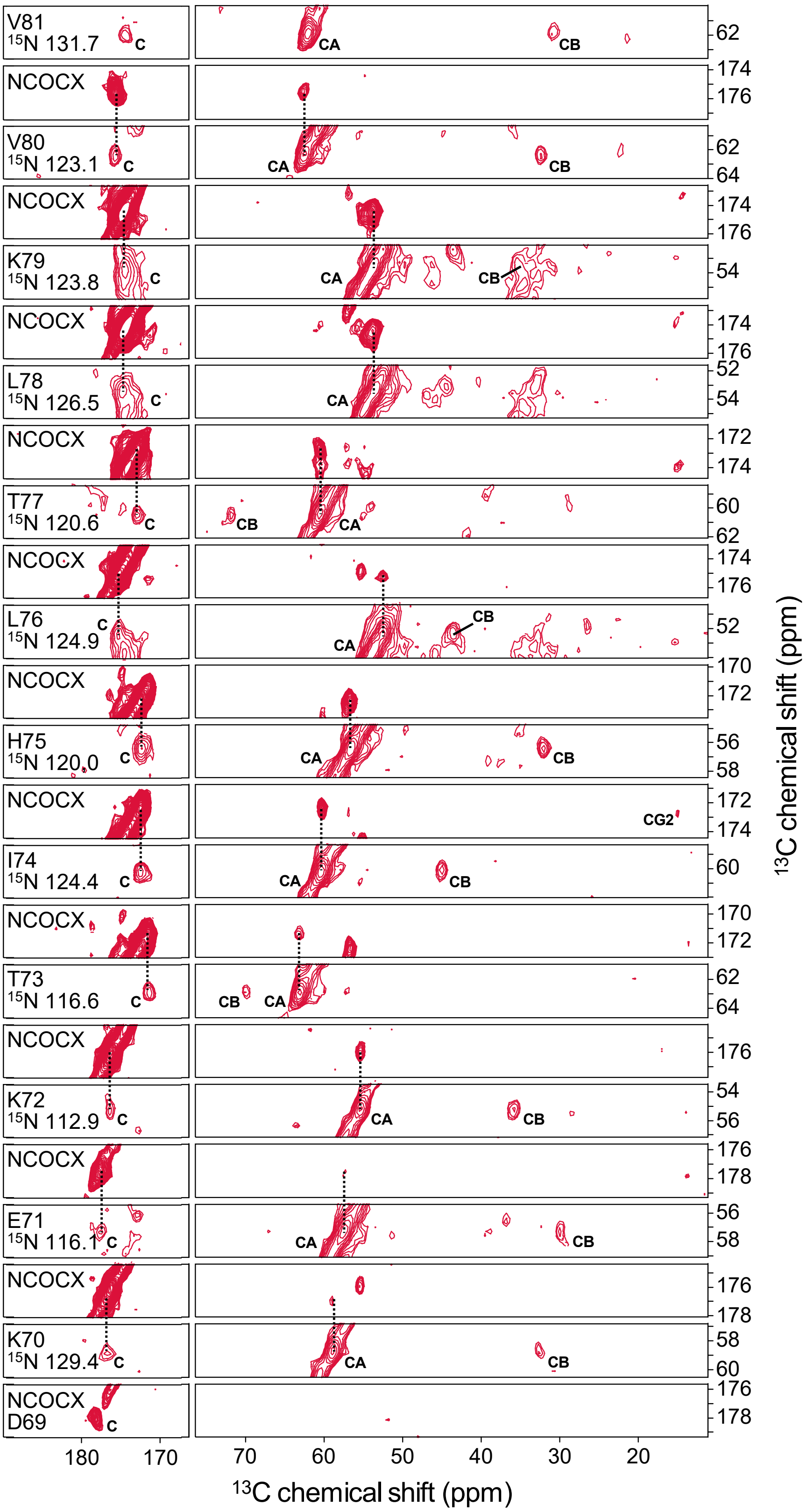

***Figure S4:*** *$^{15}$N-$^{13}$C-$^{13}$C NCOCX/NCACX strip plot showing the backbone walk from residues D69 to V81. The N nuclei of I68 and D69 are not observed, but, just like for the complex of N-FAT10-C0 with NUB1L, proximity of these two residues was confirmed by $C_{\gamma2}$-$C_{\alpha}$ and $C_{\gamma2}$-$C_{\beta}$ cross peaks in DARR spectra with long mixing times.*

### Chemical shift assignments for N-FAT10-WT in complex with NUB1L

***Table S3:*** *$^{15}N$ and $^{13}C$ chemical shifts (ppm) of U-$^{13}C$,$^{15}N$-N-FAT10-WT (amino acids 5-86) co-sedimented with natural abundance NUB1L.*

| Residue | N | C' | Cα | Cβ | Cγ | Cδ | Other |
|---|---|---|---|---|---|---|---|
| A5 | 40.1 | 173.6 | 52.0 | 20.4 | | | |
| W17 | | | | | | | Cε2: 138.9, Nε1: 129.9 |
| I68 | | 174.3 | 60.2 | 39.4 | 28.0, 16.8 | 13.7 | |
| D69 | | 178.0 | 52.6 | 40.9 | 179.2 | | |
| K70 | 129.4 | 176.8 | 58.9 | 32.8 | 24.9 | 29.4 | Cε: 42.0, Nζ: 32.7 |
| E71 | 116.1 | 177.6 | 57.5 | 29.8 | 36.6 | 183.8 | |
| K72 | 112.9 | 176.2 | 55.3 | 35.8 | 25.1 | 29.2 | Cε: 42.0, Nζ: 32.7 |
| T73 | 116.6 | 171.5 | 63.1 | 69.9 | 22.4 | | |
| I74 | 124.4 | 172.5 | 60.4 | 45.0 | 31.2, 14.9 | 16.8 | |
| H75 | 120.0 | 172.4 | 56.5 | 32.2 | 129.5 | 120.2 | Cε1: 134.6, Nδ1:182.2, Nε2: 175.1 |
| L76 | 124.9 | 175.2 | 52.6 | 43.5 | 27.0 | 23.1 | |
| T77 | 120.6 | 172.9 | 60.5 | 71.9 | 22.1 | | |
| L78 | 126.5 | 174.7 | 53.5 | 43.9 | 26.9 | | |
| K79 | 123.8 | 174.6 | 53.7 | 35.3 | 24.0 | 29.4 | Cε: 41.9, Nζ: 32.7 |
| V80 | 123.1 | 175.6 | 62.6 | 32.4 | 22.1,21.2 | | |
| V81 | 131.7 | 174.4 | 62.1 | 30.9 | 21.2 | | |
| E | | 181.2 | 58.1 | 31.2 | 36.6 | 184.5 | |
| I | | | 61.0 | 38.5 | 27.5, 17.6 | 13.1 | |
| L | | 174.7 | 54.0 | 45.9 | 27.9 | | |
| P | 135.2 | 176.6 | 63.5 | 32.3 | 27.5 | 50.6 | |
| R | | | | | | 43.4 | Cζ: 159.5, Nε: 84.3, Nη: 71.6 |
| S | 113.9 | 174.4 | 58.3 | 63.9 | | | |
| S | | 173.2 | 57.1 | 66.1 | | | |
| V | 126.5 | 174.1 | 60.5 | 35.9 | 21.5 | | |
| V | 120.9 | | 66.2 | 31.5 | | | |
| V | 120.8 | 177.7 | 66.8 | 31.6 | | | |
| Y | | | | | | | Cζ: 158.1 |

### 6. Torsion angle predictions for N-FAT10-WT in complex with NUB1L

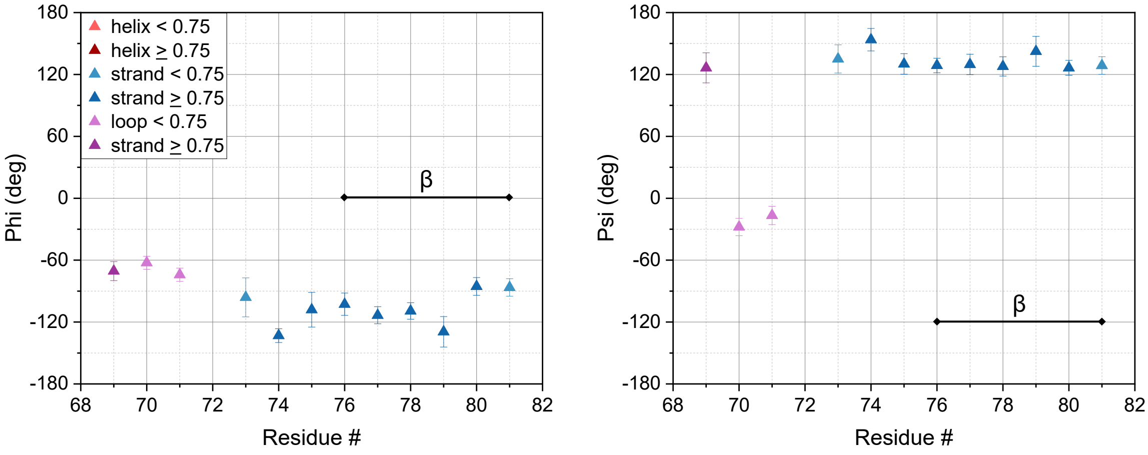

***Figure S5:*** *Backbone torsion angle and secondary structure prediction based on MAS NMR of N-FAT10-WT co-sedimented with NUB1L. Error bars correspond to the standard deviations of the Φ and Ψ angles of the best matches in the TALOS-N database. For residue K72, there is no consensus among the database matches and, hence, no predicted torsion angles are shown. Secondary structure predictions are based on observed chemical shifts and classified as helix, strand, or loop. Predictions with a confidence ≥ 0.75 are highly reliable. Based on sequence information, the secondary structure of I68 is classified as loop. Based on observed chemical shifts, the secondary structure of K72 is classified as loop.*
